# Genetic ablation of visual perception reveals behaviour changes in male and female malaria mosquitoes

**DOI:** 10.64898/2026.03.03.709228

**Authors:** Dennis Klug

## Abstract

The role of vision in the behavior of blood-feeding mosquitoes has remained largely overlooked, particularly in species of the *Anopheles* genus, despite their significant impact on global health. While the importance of olfactory and thermal cues in host-seeking is well established, the contribution of visual stimuli to mating and feeding behavior remains far less understood. In particular, *Anopheles* mosquitoes exhibit complex swarming behavior that depends on visual input, suggesting an underexplored avenue of research with direct implications for vector control. This study introduces a genetically modified mosquito line lacking the enzyme Tan, a hydrolase involved in both dopamine and histamine metabolism, to investigate the behavioral relevance of visual cues in *Anopheles*. Through a combination of behavioral assays and controlled laboratory experiments, the impact of visual disruption on attraction to UV-B light, host-seeking, and blood-feeding success was assessed. The findings demonstrate a reduced light-dependent attraction in both *Anopheles* males and females, suggesting an impairment in visual processing or a related behavioral response. This observation has implications for reproductive success and potential adaptation to anthropogenic environments with artificial light. By leveraging this novel knockout model, the study offers new tools and perspectives to better understand how vision shapes mosquito behavior and how this knowledge could be harnessed in the development of next-generation vector control strategies.

## Introduction

Female mosquitoes require a blood meal to invest in reproduction and lay eggs. Therefore, efficient host-seeking is a critical factor for species survival. To locate a suitable host, female mosquitoes rely on specific cues, particularly heat and carbon dioxide (CO₂) (Montell, 2025). However, additional signals—such as aldehydes found in human sweat—are also detected and integrated to enhance host-seeking (Zhao *et al*., 2022). More recently, it has been demonstrated that female *Aedes* mosquitoes utilize visual cues to identify potential hosts from a distance before cues like smell and heat can be detected (van Breugel *et al*., 2015; Hawkes and Gibson, 2016; Chandel *et al*., 2024). In addition, it was shown that mosquitoes use certain colors to identify hosts suitable for blood feeding (Alonso San Alberto *et al*., 2022). The subsequent integration of all these diverse cues in the mosquito brain is crucial for rapid and species-specific host seeking and, as a result, for the success of blood feeding (Zhao *et al*., 2022; Montell, 2025).

Much of what we know about host-seeking behavior in blood-feeding insects stems from studies on *Aedes* species, particularly *Aedes aegypti* (Montell, 2025). This preference can be partially attributed to practical considerations: *Aedes* eggs can be stored in a dried state for up to two years, greatly reducing the labor required for continuous colony maintenance. Another contributing factor to the popularity of *Aedes* mosquitoes in behavioral research is their diurnal activity. Female *Aedes* are active during the day, seeking hosts both indoors and outdoors, with peak activity shortly after sunrise and just before sunset. In contrast, malaria mosquitoes from the genus *Anopheles* predominantly blood-feed at night—particularly between 10 p.m. and 4 a.m.—which may explain the prevailing assumption that visual cues are less relevant or even negligible for these species.

In this study, I investigated a loss-of-function mutant of the hydrolase Tan in the malaria mosquito *Anopheles coluzzii*. Tan is a dual-function enzyme that converts N-β-Alanyldopamine (NBAD) to dopamine in both the cuticle and central nervous system (Massey and Wittkopp, 2016), and also catalyzes the conversion of carcinine to histamine in the retina of insects and crustaceans (**Fig. 1**) (Borycz *et al*., 2002). NBAD and dopamine are precursors for various pigments incorporated into the cuticle, while histamine serves as a key neurotransmitter in the retina.

**Figure 1.**
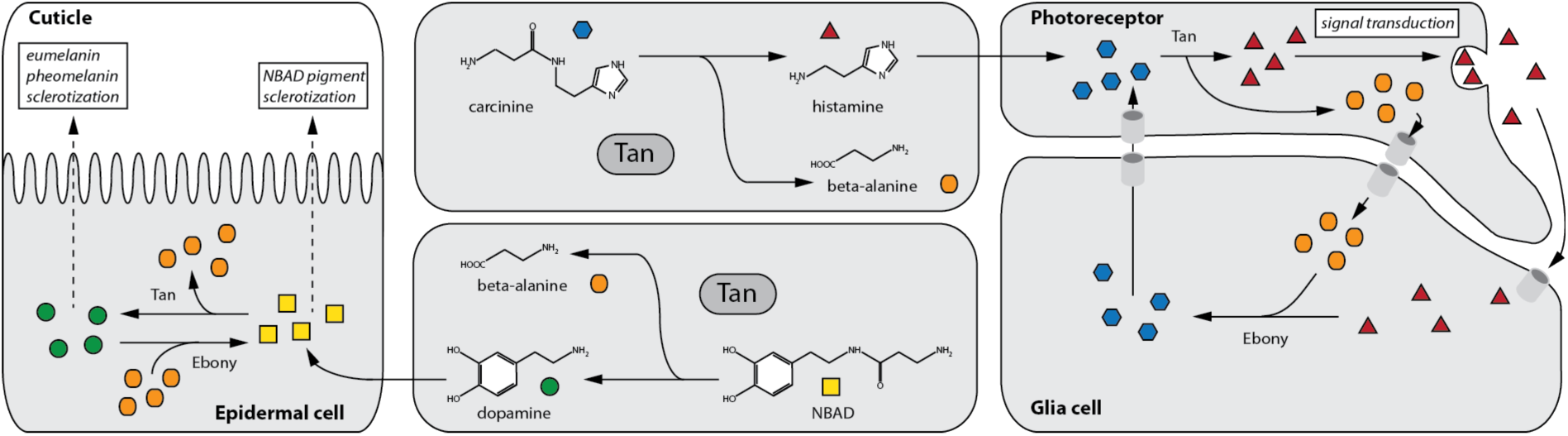
The *tan* gene encodes a hydrolase with two distinct functions. Depending on the tissue, Tan catalyzes two distinct chemical reactions. In epidermal cells, Tan hydrolyzes N-β-alanyl dopamine (NBAD) into β-alanine and dopamine. Subsequently, dopamine or its derivatives are secreted into the cuticle as precursors for pigments (e.g., eumelanin, pheomelanin, and compounds required for sclerotization). Although NBAD itself also serves as a pigment precursor, the absence of Tan alters the ratio between dopamine and NBAD, leading to changes in the pigment composition of the cuticle. In photoreceptors, Tan is required to replenish the neurotransmitter histamine from carcinine. Carcinine, synthesized in glial cells, is taken up by photoreceptors and converted by Tan into histamine and β-alanine. During signal transmission, histamine is released into the synaptic cleft, from where it can be reabsorbed by glial cells. In glial cells, the NBAD-synthase Ebony regenerates carcinine from histamine and β-alanine.

By catalyzing these two reactions, Tan influences cuticle coloration by regulating the availability of NBAD and dopamine and ensures the continuous transmission of visual signals from the retina to the brain by recycling histamine from carcinine. In contrast, the enzyme Ebony, a β-alanyl biogenic amine synthetase, uses β-alanine to convert dopamine and histamine into NBAD and carcinine, respectively (**Fig. 1**) (Borycz *et al*., 2002; Richardt *et al*., 2003). Together, Tan and Ebony represent a reciprocal enzyme pair maintaining homeostasis in cuticle pigmentation and visual neurotransmission.

Beyond its unique enzymatic duality, Tan has interesting structural features. It shares remarkable similarity with isopenicillin-N-acyltransferase (IAT) from *Penicillium chrysogenum*, which catalyzes the final step in penicillin biosynthesis by transferring an acyl group to isopenicillin N (Bokhove *et al*., 2010). Due to the high degree of structural similarity (**Fig. 2A**), it has been speculated that Tan may have an ancestral origin (True *et al*., 2005). Interestingly, Tan is highly conserved among insects and crustaceans but absent in the other two major arthropod subgroups—myriapods and chelicerates (**Fig. 2B**). While this pattern suggests vertical inheritance, it also raises the possibility of horizontal gene transfer in a common ancestor of insects and crustaceans. Structurally, both IAT and Tan are composed of a small and a large subunit, which are connected by a highly conserved C–G cleavage site (**Fig. 2C, E**). Proteolytic cleavage of the proprotein is required to activate IAT, thereby releasing the large subunit that contains the catalytically active peptidase C45 domain (**Fig. 2C**). This cleavage site is also conserved in Tan which suggests proteolytic processing, this has so far been not shown experimentally. Across various insect species (*A. aegypti*, *A, gambiae* and *D. melanogaster*) and the crustacean *Penaeus vannamei*, Tan exhibits a high degree of sequence conservation, with only few regions showing greater variability (**Fig. 2E, Fig. S1**).

**Figure 2.**
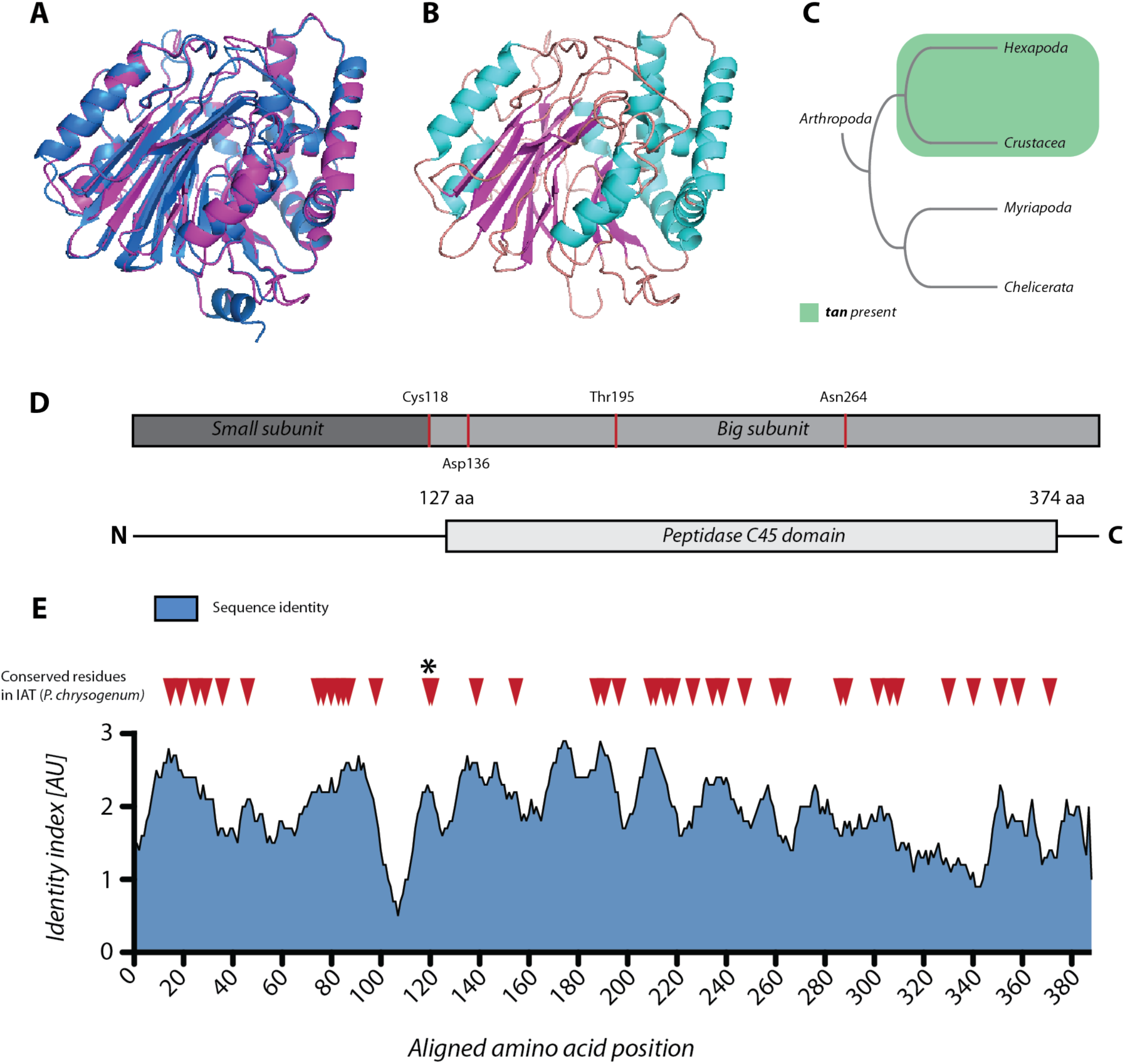
Tan is closely related to fungal isopenicillin-N-acyltransferases (IATs) and conserved in hexapoda and crustaceans. **A)** Superposition of modelled AgTAN (purple) with the crystal structure of the IAT from *Penicillium chrysogenes* (blue) (PDB: 2X1C). **B)** Modelled tertiary sturcture of AgTAN, ß-strands are shown in purple while α-helices are depicted in turquoise. **C)** Simplified phylogenetic tree of the arthropda. TAN distribution is restricted to insects and crustaceans (shaded in light green). **D)** Domain organization of TAN. TAN is composed of a small and a big subunit which mature through cleavage between G117 and C118 (*Ag*TAN) from a pre-protein. The IAT-specific peptidase C45 domain localizes in the big subunit between V123 and Y369 (*Ag*TAN). Residues predicted to form the catalytic center (Cys118, Asp136, Thr195 and Asn264 in *Ag*TAN) are indicated as red stripes. **E)** Conservation of TAN. Shown is the sequence identity between the TAN homologues *Dm*TAN (*Drosophila melanogaster*), *Aa*TAN (*Aedes agypti*) and *Ag*TAN (*Anopheles gambiae*) as well as the homologue from the whiteleged shrimp *Penaeus vannamei Pv*TAN. Residues which are also conserved in *Pc*IAT are indicated as red arrowheads. The highly conserved G-C cleavage site is indicated with a black asterisk. The dataset was smoothed using a 10-position running average.

Due to its role in cuticle pigmentation, Tan has been studied extensively in the context of transcriptional regulation and speciation in *Drosophila* (Jeong *et al*., 2008; Wittkopp *et al*., 2009), while its function in the retina has provided insights into insect visual physiology (Borycz *et al*., 2002; True *et al*., 2005). According to the proposed mechanism, Tan null mutants are thought to suffer from impaired vision, as histamine levels in the retina are reduced due to the absence of histamine recycling. However, *de novo* synthesis of histamine from other sources may compensate under dark conditions or help maintain a minimal level of visual signal transmission during daylight. This raises the question of how much visual information **Tan** null mutants can still perceive.

Physiologically, *Drosophila tan* mutants show significantly reduced histamine concentrations in the head and exhibit abnormal electroretinogram (ERG) responses (True *et al*., 2005). However, to the best of my knowledge, behavioral studies directly assessing visual performance in tan null mutants are currently lacking.

In this study, I used a *tan* null mutant to investigate visual perception in *Anopheles* mosquitoes by quantifying their attraction to UV-B light and assessing host-seeking efficiency and blood feeding success under daylight conditions.

## Results

### *Tan(-)* mosquitoes show a reduced likelihood of being caught by UV-B-based mosquito traps

Loss of *tan* expression in *Drosophila melanogaster* has been shown to disrupt histamine recycling in the visual system, resulting in abnormal electroretinogram (ERG) responses (True *et al*., 2005). Therefore, I hypothesized that *tan(-)* mosquitoes may exhibit impaired vision, potentially reducing their susceptibility to light-based mosquito traps. To test this hypothesis, a loss-of-function mutant was generated through the knock-in of a cassette conferring resistance to the antibiotic puromycine which replaced the first exon of *tan* including the start codon (**Fig. S2A**). Following transgenesis, integration of the cassette was confirmed by PCR and absence of transcription of *tan gene* was verified by RT-PCR (**Fig. S2B,C**). Phenotypically, *tan(-)* mosquitoes displayed a general brightening of the cuticle which was in particular visible in the abdomen by comparing areas usually highly pigmented in the wild-type (**Fig. S2D**). To test a visual impairment of *tan(-)* mosquitoes, I developed an experimental setup using a large mosquito cage (324 L) and a commercially available UV-B-based mosquito trap (**Fig. S4A, B; Fig. 3A**). The trap includes an internal fan that creates negative pressure, drawing in mosquitoes that approach the opening and causing death by dehydration. Two sugar feeders were placed beside the trap to provide nutrition throughout the experiment (**Fig. S4B, C**). In initial experiments, equal numbers of wild-type and *tan(-)* mosquitoes of both sexes were released into the cage at 3 pm. Because the two genotypes lack easily distinguishable visual markers, mosquitoes were dusted with fluorescent powder (Moonglow) following standard protocols to allow genotype identification (**Fig. 3B**; Verhulst, Loonen & Takken, 2013). After a two-hour acclimatization period (until 5 pm), the UV-B trap was activated. Trapped mosquitoes were counted the following day at 9 am. Given that night conditions began at 4 pm and ended at 2 am, the trap was active throughout the dark phase, and counting was done in daylight (**Fig. 3B**). Across five replicates, an average of 81% (± 15% SD) of wild-type mosquitoes were caught, while significantly fewer *tan(-)* mosquitoes (59% ± 24% SD) were recovered from the trap (**Fig. 3C**). The same tendency was observed in both male and female *tan(-)* mosquitoes albeit not significant for a single sex. Although these results support the hypothesis that *tan(-)* mosquitoes are less responsive to light-based traps, the long duration of the experiment (16 hours) may have allowed some mosquitoes to be caught by chance, regardless of visual attraction. To address this, the experiment was modified: the start time was moved to 1 pm, with a two-hour acclimation, and the trap was activated earlier at 3 pm. This allowed for an initial collection after two hours of trap activity (5 pm), with a final collection at 9 am the next morning. All other conditions remained unchanged. In this modified setup, *tan(-)* mosquitoes were again significantly less likely to be caught than controls (56% ± 23% SD vs. 85% ± 11% SD; **Fig. 3D**). However, this difference was only observed in the total number of collected mosquitoes; no significant difference was seen for the early collection at 5 pm. To confirm that the observed reduction in trap susceptibility was due to visual impairment—and not a general behavioral difference—the experiment was repeated with the trap’s light source covered by dark tape to block UV-B emission. Under these conditions, no significant difference in trapping rates between wild-type and *tan(-)* mosquitoes was observed (**Fig. 3E**). Although the results were consistent, I considered the possibility that fluorescent dusting itself could influence mosquito behavior. Supporting this concern, previous work has shown that dusting can reduce mosquito lifespan (Verhulst, Loonen & Takken, 2013). To eliminate this potential confounder, I planned to use *yellow(-)\** as an alternative phenotypic marker. Loss of *yellow* expression results in the absence of black pigment, giving mosquitoes a yellowish appearance due to the dominance of brown/yellow pigments (Klug et al., 2021). To test the suitability of *yellow(-)* mosquitoes for behavioral assays, I first compared their responsiveness to UV-B light with that of wild-type mosquitoes using the same setup. Catching rates for *yellow(-)* and wild-type mosquitoes were comparable (88% ± 6% SD vs. 81% ± 14% SD; **Fig. 4A**). When the trap’s light source was blocked, both groups showed similarly lower trapping rates (**Fig. 4B**), indicating that *yellow(-)* mosquitoes do not differ in visual behavior. Next, I generated a *yellow(-);tan(-)* double mutant line through genetic crosses (**Fig. S3A**), followed by enrichment via flow cytometry and puromycin selection until a homozygous colony was established which was verified through genotyping by PCR of founding females (**Fig. S3B**). *Yellow(-);tan(-)* appeared generally brighter, surpassing the phenotype of *yellow(-)* mosquitoes (**Fig. S3C**). *Yellow(-);tan(-)* mosquitoes were subsequently tested against wild-type mosquitoes in the UV-B trap assay. Mosquitoes lacking expression of *tan* and *yellow* showed significantly lower trapping rates than wild-type which was even significant if only males were considered (49% ± 13% SD vs. 78% ± 15% SD; **Fig. 4C**). Surprisingly, this difference persisted even when the UV-B light was blocked (**Fig. 4D**), suggesting that the absence of *yellow* may enhance the *tan(-)* phenotype.

**Figure 3.**
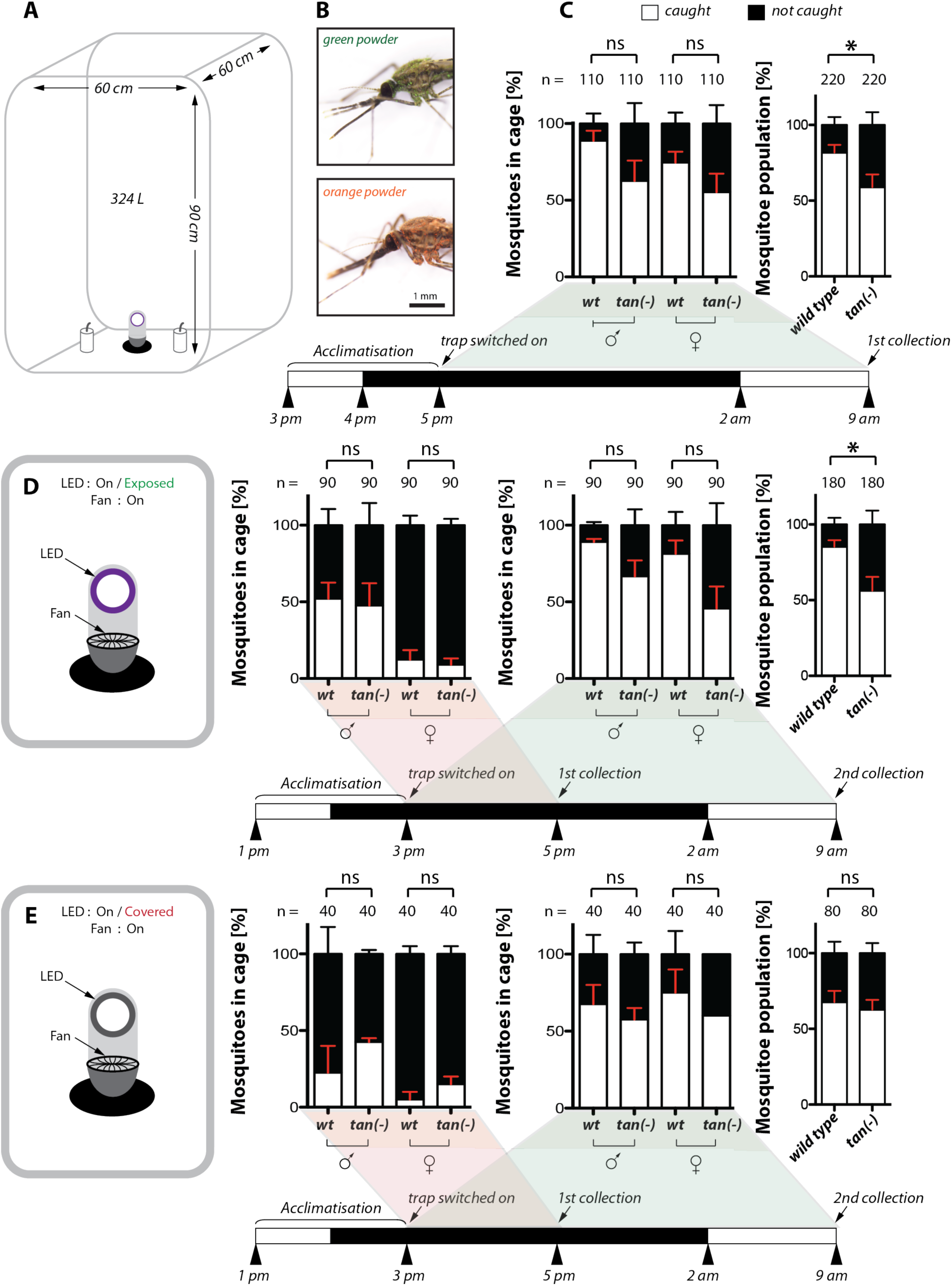
Tan deficient mosquitoes are less attracted by UV-B light. **A)** Experimental setup to evaluate attraction of mosquitoes to UV-B light. A cage with 324 L volume (90 cm x 60 cm x 60 cm) was set up with a commercial mosquito light trap. Two sugar dispensers were placed in the shade of the light trap to avoid the catch of feeding mosquitoes. **B)** Mosquitoes were marked with orange and green fluorescent powder (Moon Glow) to differentiate the genotype of caught mosquitoes. Scale bar: 1 mm. **C)** Percentages of caught and not caught mosquitoes according to sex and genotype with one collection timepoint. The graph on the right summarizes the data displayed on the left independently of sex. Shown data represent five independent experiments and were tested for significance with the Mann Whitney test. p value = 0.0353*. ns: not significant. **D)** Repetition of the experiment depicted in (C) with a modified collection regime (two timepoints). Pictogram on the left depicts the state of the light trap (fan switched on, LEDs uncovered). The duration of the experimental setup was extended by two hours and a first collection of caught mosquitoes was performed after two hours. The light trap was switched on for the whole course of the experiment and LEDs were uncovered. The graph on the left represents the ratio of caught and not caught mosquitoes after two hours (red shade). The graph in the middle summarizes the total percentage of caught and surviving mosquitoes according to sex (green shade). The graph on the right displays the same data according to genotype while males and females are represented in the same column. Shown data represent three experiments and were tested for significance with Mann Whitney test. p value = 0.0185*. ns: not significant. **E)** Repetition of the experiment depicted in (D) with two collection timepoints but covered LEDs. Pictogram on the left depicts the state of the light trap (fan switched on, LEDs covered). Data are presented the same way as in (D). Shown data represent two experiments and were tested for significance with Mann Whitney test. ns: not significant.

**Figure 4.**
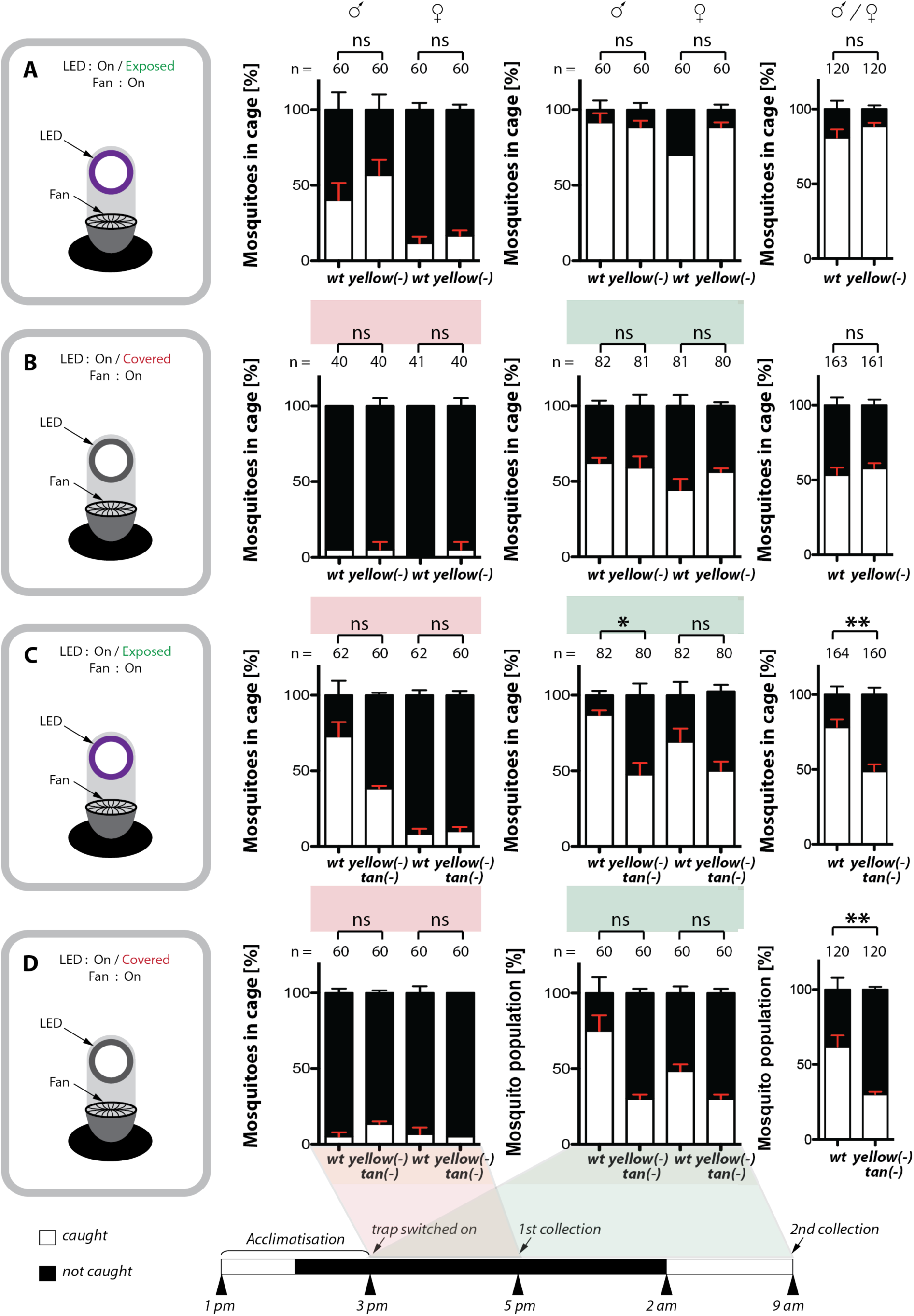
The use of *yellow\** as a phenotypic marker confirms the impact of *tan(-)* on behaviour experiments. **A**) Percentages of caught and not caught mosquitoes according to sex and genotype with two collection timepoints. *The yellow(-)* lines was tested in comparison to control mosquitoes. The graph on the left represents the ratio of caught and not caught mosquitoes after 2 hours. The graph in the middle summarizes sexp-specific data for total caught and not caught mosquitoes and the graph on the right displays data of total collected mosquitoes independent of sex. Shown data represent three biological replicates. Statistics were performed with Mann Whitney test. ns: not significant. The illustration at the bottom inidcates the day/night cycle mosquitoes in (A), (B), (C) and (D) were exposed to during the experiment as well as acclimatization phase and collection time points. **B)** Repetition of the experimental setting as in (A) with covered light source. Shown data represent three biological replicates. Statistics were performed with Mann Whitney test. ns: not significant. **C)** Testing of *yellow(-):tan(-)* in competition with control males and females. Shown data represent three independent experiments. Statistics were performed with Mann Whitney test. p value = 0.0294* (males), p value = 0.0071** (both sexes combined), ns: not significant. **D)** Repetition of the experimental setting as in (C) with covered light source. Shown data represent three biological replicates. Statistics were performed with Mann Whitney test. p value = 0.0048**, ns: not significant.

### Loss of *tan* does not affect blood-feeding success of females at short range

Next, I investigated whether the impaired visual perception observed in *tan(-)* mosquitoes affects the blood-feeding success of females. In *Aedes* mosquitoes, visual cues have been shown to play a key role in host detection (van Breugel *et al*., 2015; Chandel *et al*., 2024). For each experiment, 20 females were placed into the experimental cage and allowed to acclimate for at least 2 hours under daylight conditions (**Fig. 5A**). After acclimatization, females were given a 15-minute window to blood-feed on my forearm. On average, 45% (± 9% SD) of wild-type females successfully obtained a blood meal within this period (**Fig. 5B**). Visual inspection indicated that most fed females were fully engorged, with minimal variation in blood bolus size, suggesting that feeding generally began within the first 5 minutes (**Fig. 5C, D**). Comparable feeding rates were observed for the *yellow(-)* phenotypic marker line (41%, ± 16% SD), and even slightly higher but not significantly different feeding rates (60%, ± 16% SD) were recorded in *yellow(-);tan(-)* females, indicating that the absence of *Tan* does not impair short-range host seeking or blood-feeding behavior (**Fig. 5B**).

**Figure 5.**
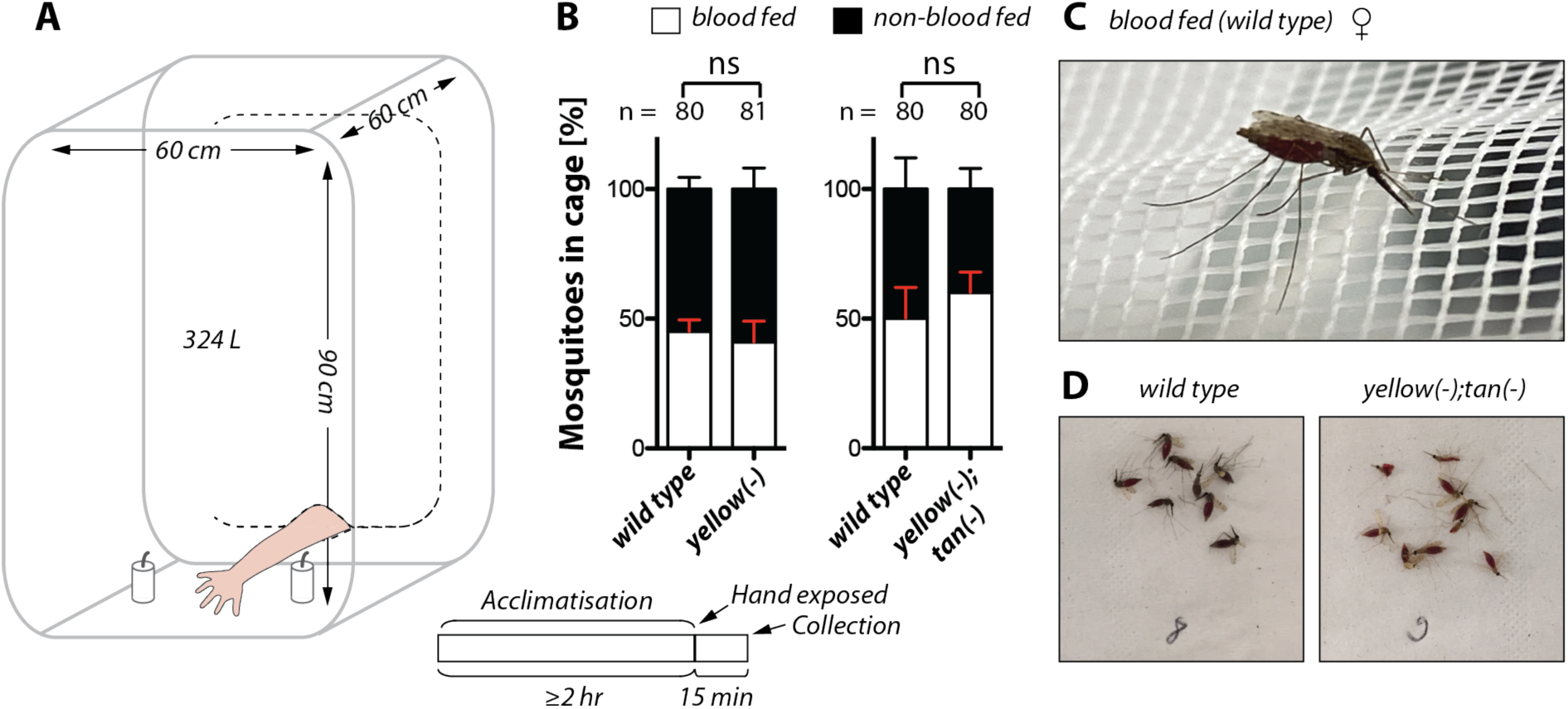
*Tan(-)* mosquitoes display normal host seeking and blood feeding behaviour at short distances. **A)** Experimental setup to evaluate host seeking behaviour and blood feeding. A cage with 324 L volume (90 cm x 60 cm x 60 cm) was set up with 20 females of each genotype and allowed to acclimatize for ≥2 hours. Subsequently the hand of the experimentator was exposed within the cage to attract mosquitoes and allow blood feeding. After 15 minutes mosquitoes were evaluated for the presence of a blood bolus within the abdomen. **B)** Ratio of blood fed and non fed females. Wild-type females were tested in competition with *yellow(-)* or *yellow(-);tan(-)* females. Data represent 4 biological replicates and were tested with a Mann Whitney test to determine statistical significance. ns: not significant. **C)** Example image of a blood fed wild-type female. **D)** Separation and counting of blood fed wild-type and *yellow(-);tan(-)* females after completion of an experiment.

### *Tan(-)* mosquitoes show no transgenerational fitness costs but exhibit reduced lifespan

Visual cues are known to influence swarming behavior in *Anopheles coluzzii*, supporting collision avoidance and mate location (Gupta *et al*., 2024). To assess whether visual impairment in *tan(-)* mosquitoes affects mating and reproduction, I conducted a long-term cage experiment using a mixed colony containing equal proportions of wild-type and homozygous *tan(-)* males and females. To enable sex-specific tracking, wild-type and *tan(-)* males carried a GFP-marked Y chromosome (Bernardini *et al*., 2014)(**Fig. 6A**). In each generation, offspring were sorted via flow cytometry into GFP-positive (males) and GFP-negative (females) larvae (**Fig. 6B**). Larvae were counted and subjected to puromycin treatment following standard protocols (Volohonsky et al., 2015) (**Fig. 6A**). By comparing initial counts to post-selection survivors, the proportion of *tan(-)* carriers in each sex was determined (**Fig. 6C**). Across ten generations, the genotype frequencies stabilized around the expected Hardy-Weinberg equilibrium for X-linked inheritance, indicating no significant transgenerational fitness costs under laboratory conditions (**Fig. 6C**). Despite the absence of detectable fitness effects in mixed populations, I also evaluated the longevity of *tan(-)* females compared to controls. From day 10 post-hatching onward, *tan(-)* females showed a significantly increased mortality rate (**Fig. 6D**). On average, wild-type females lived up to 10 days longer than their *tan(-)* counterparts, suggesting that loss of *Tan* hydrolase activity may have systemic physiological consequences.

**Figure 6.**
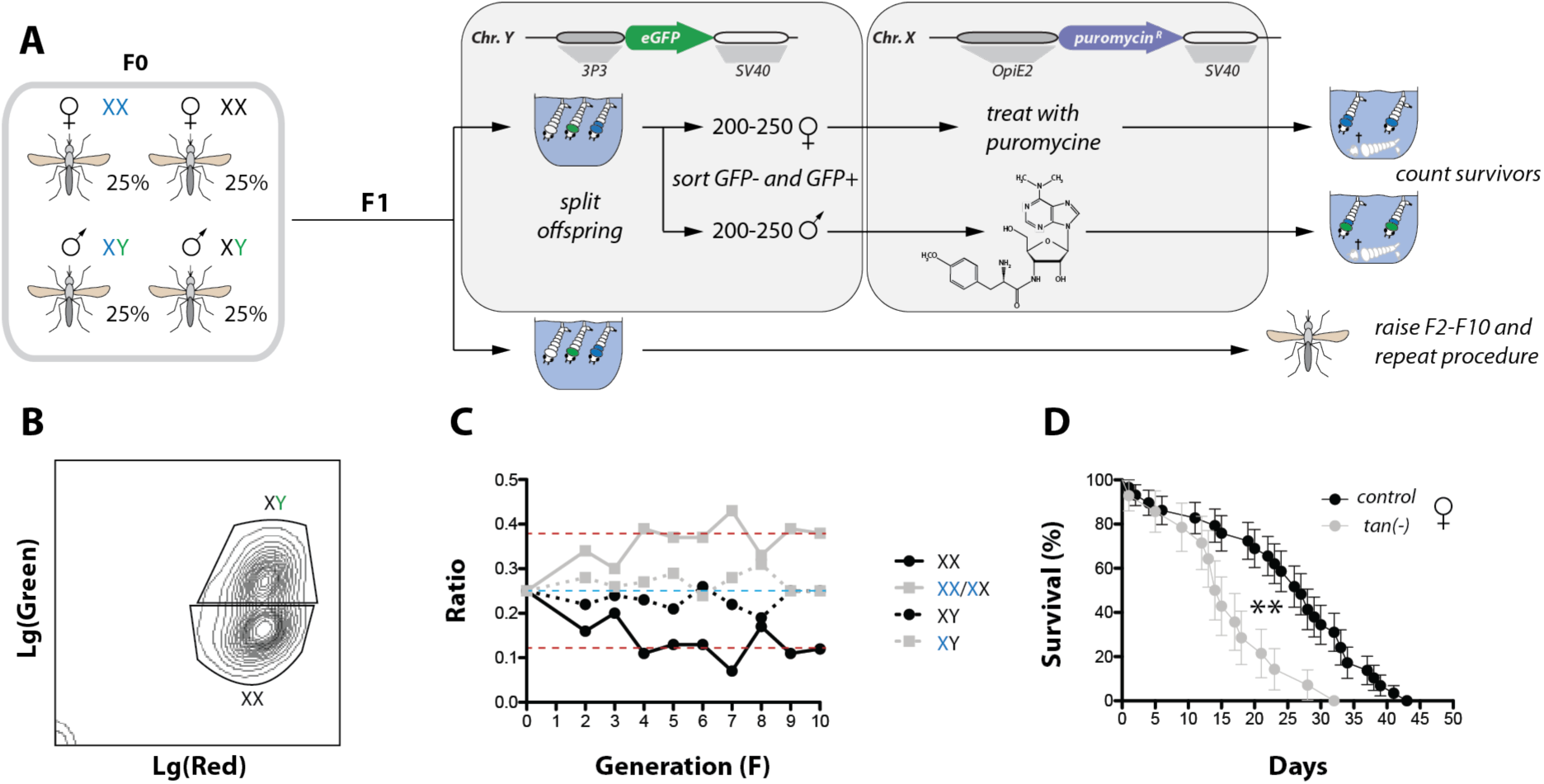
*Tan(-)* mosquitoes display no transgenerational fitness costs but have a reduced lifespan. **A)** The fitness of the *tan(-)* line was assessed in competition with the respective control (*cas9::RFP+*). To allow sex sorting mosquitoes were crossed with the TS1 line expressing GFP on the Y chromosome. In the founder population 50 *tan(-)* males positive for GFP were mixed with 50 *tan(-)* females, 50 control males positive for GFP and 50 control females. The F1 offpring was divided in two populations. One half of the population was used to propagate the colony while the other half was used for sex sorting using COPAS. Of this population 200-250 males and 200-250 females were sorted and treated with puromycine. After three days the number of survivors was counted to calculate the presence of the *tan(-)* transgene in the population of the respective sex. This protocol was repeated each generation until F10. **B)** Gating strategy to separate male and female mosquito larvae. Larvae form two distinct populations based on the presence or absence of male specific GFP expression. Note that both populations display a red shift because all larvae are homozygous for *cas9:rfp+*. **C)** Presence of the *tan(-)* transgene in competition with the wild-type allele during consecutive breeding for 10 generations. Red dotted lines indicate the Hardy-Weinberg equilibrium for control and heterozygous/homozygous *tan(-)* females and the blue dotted line for wild-type/hemizygous males. **D)** Life span measurements of female mosquitoes. Females homozygous for *tan(-)* and respective control females were kept on 10% sucrose solution and the number of dead mosquitoes per day was monitored over time. Data represent two independent experiments with >30 mosquitoes each and were tested for significance with the Log-rank (Mantel-Cox) test. p value = 0.0016**.

## Discussion

Visual perception in blood-feeding insects remains a largely understudied area and has been investigated primarily in *Aedes* mosquitoes. This is somewhat surprising, given that malaria-transmitting mosquitoes of the genus *Anopheles* are still responsible for over 500,000 deaths each year—an order of magnitude higher than the approximately 10,000 deaths caused by *Aedes*-borne dengue (WHO, 2024).

My characterization of the *tan(-)* mutant’s response to artificial light provides a genetic tool that may, in future studies under more naturalistic conditions, help elucidate the role of vision in behaviors such as swarming and host-seeking. Such knowledge could ultimately contribute to the development of novel intervention strategies aimed at disrupting blood feeding and reproduction in *Anopheles* mosquitoes. Indeed, it has recently been shown that especially male *Anopheles* mosquitoes use visual cues within swarms to avoid collisions and track females (Gupta *et al*., 2024). However, the absence of a genetic model with reduced eyesight limits the interpretation of these results. *Tan(-)* mosquitoes could possibly fill this gap and serve as a valuable tool to decipher the effect of vision on behaviour in *Anopheles*. The use of a high-intensity UV-B stimulus provided a robust and reproducible method to initially characterize the behavioral phenotype of the *tan(-)* mutant under controlled laboratory conditions. Loss of Tan significantly decreased attraction of males to UV-B light based mosquito traps by ∼36% (mean) compared to control males (**Fig. S5**). Similarly, females also showed a signficant effect with a reduction of ∼35% (mean) albeit catching rates for males were generally higher compared to females (∼16% difference between males and females for wild-type) (**Fig. S5**). This effect was consistent across all experiments regardless of whether mosquitoes were marked ectopically with fluorescent powder or phenotypically through the additional knockout of yellow. Covering of the light source cancelled out any effect on the catching rate (**Fig. 3E**). This result strongly indicates that the observed phenotype is light-dependent and not an artifact of other trap features like airflow. The only exception to this observation was the *tan(-):yellow(-)* double knockout were low catching rates were preserved even when the light source was covered while *tan(-)* and *yellow(-)* single mutants displayed no difference in the catching rate to controls within this experimental setting. This observation could be explained by different reasons. I used only commercially available equipment in this study, therefore the mosquito trap was not designed for behaviour experiments why light and fan couldn’t be controlled independently. As a consequence the light was covered with black tape which greatly reduced light emission but was not absolute, possibly leaving a weak stimulus for UV-B light. Another effect could be attributed to the phenotypical marker *yellow\**. Yellow is required for the synthesis of black eumelanin in all tissues of insects. In the absence of Yellow eumelanin is missing making mutants appear yellow because of the remaining brown and yellow pigments. It is possible that the absence of black pigment in the ommatidia of the mosquito itself has an effect on vision, even if this was not detectable in the *yellow(-)* single mutant. This could have possibly enhanced the visual defects in *tan(-)* mosquitoes leading to a more penetrating phenotype. Indeed, it has been shown that tan and yellow act in concert during pigment formation (Jeong *et al*., 2008).

In competition with wild-type the *tan(-)* transgene displayed no transgenerational fitness costs over the course of 10 generations. This was somehow suprising since vision has been described to be an important cue for *Anopheles* males to find females which would suggest a reduction in mating success (Gupta *et al*., 2024). However, the breeding conditions in the laboratory are vastly different compared to nature. Mosquito colonies in climate chambers are usually kept in relatively small cages which do not allow any swarm formation possibly making visual cues less important. Similarly, I was not able to detect changes in blood feeding success of *tan(-)* females. This could simply be because the initial distance between host and female mosquitoes was to short making heat, CO_2_ and odor plumes the dominant cues which negated the impact of an impaired vision. For *Aedes* it has been described that visual cues are important to identify potential hosts only from further away (van Breugel *et al*., 2015). It would be interesting to test the behaviour of *tan(-)* males females under conditions closer mimicking the natural environment. Previous observations indicate that visual cues are important to guide the behaviour of *Anopheles* males in mating swarms (Gupta *et al*., 2024). Indeed, males displayed a very high catching rate in UV-B based trap assasy (∼88% of wild-type males had been caught after 18 h of trap exposure) but impairing vision through the loss of Tan led to a similar reduction of catches in both sexes indicating that vision is also important for females (**Fig. S5**). This would indirectly imply a role for Tan in mating success and possibly, in host seeking, both important parameters for fitness and species survival. This assigns Tan an important role in the adaption of *Anopheles* mosquitoes to urbanized areas with high light pollution. In these environments tightly controlled tissue-and time-specific expression of Tan could be an important driver of evolution by modifying the behaviour of mosquitoes to become less distracted by lamps, lanterns or even, like in this study, light-based mosquito traps. Such selected mosquitoes would be more likely to engage in reproduction possibly driving speciation and the adaption to humans as a host for blood feeding. Due to the dual role of Tan in pigmentation this selective pressure could possibly even translate into phenotypic differences affecting the coloration of the cuticle as it has been shown for different *Drosophila* species (Jeong *et al*., 2008). It is important to note, however, that the experimental conditions tested during this study are artificial due to the restricted space in the insectory and the testing of UV-B light as only attractant. The relevance of these findings to natural behaviors, such as mate-finding in swarms or host-seeking by moonlight, remains to be determined.

In summary, our understanding of how visual cues influence the behavior of blood-feeding mosquitoes remains limited. Gaining deeper insights into the role of vision in host seeking and mating in *Anopheles* mosquitoes is urgently needed to inform and improve intervention strategies aimed at reducing malaria transmission.

## Materials and methods

### Bioinformatic analysis

Prediction of secondary and tertiary structure of AgTAN (AGAP000023) was performed with i-TASSER (Roy, Kucukural and Zhang, 2010) and sequences were aligned with Clustal MUSCLE (Edgar, 2004). The identity and similarity indexes were created as described previously (Klug and Frischknecht, 2017). Editing of the modelled AgTAN tertiary structure was performed with PyMOL v1.7.4.4 (The PyMOL Molecular Graphics System, Schrödinger, LLC).

### Mosquito breeding

*A. coluzzii* (genetic background: *Ngousso*) mosquitoes were kept at standard conditions (27-28°C, 75-80% humidity, 12-hr/12-hr light/dark cycle). Larvae were hatched in deionized water and fed with finely pulverized fish food (TetraMin). After pupation, pupae were collected in small glass dishes and transferred into netted cages. Hatched mosquitoes were fed with 10% sugar solution *ad libitum* using the capillarity of a cord, dipping in a sample containers filled with sugar solution. To propagate colonies, four to seven day old mosquitoes were blood fed for 10-15 minutes on anesthetized mice. Two days after feeding mosquitoes were offered a glas dish with wet filter paper to allow egg laying. L1 larvae hatched after two days and were raised as described. For light trapping experiments *tan(-)* and control mosquitoes were raised separately and mixed as adults. If experiments were performed with *yellow(-)* or *yellow(-);tan(-)* mosquitoes, neonate larvae of both lines were mixed to equal shares with controls and raised together.

### Construct cloning for site-directed mutagenesis of *tan*

The cloning of the *pENTR-Tan* plasmid used for transgensis has been performed as described previously (Klug *et al*., 2022). In brief, three guide RNAs (gRNAs) targeting the first exon of *tan* (AGAP000023) have been manually selected and tested for potential off-target sites using http://www.rgenome.net/cas-designer/ (Cas-OFFinder) (Bae, Park and Kim, 2014). The gRNAs have been ordered as single stranded oligonucleotides from IDT DNA, annealed and cloned into the *pKSB-sgRNA* plasmid (Addgene #173671–3) creating three separate gRNA building blocks. For the construction of the repair template, regions of homology on the 5’ side (DK033 and DK034) and the 3’side (DK035 and DK036) of the sequence targeted by the gRNAs were amplified from genomic wild-type (*Ngousso*) DNA and subcloned into *pJET* using the CloneJet PCR cloning kit (Thermo Fisher Scientific) to generate the two bulding blocks *pJET-5’tan* and *pJET-3’tan*. After sequence validation of all plasmids, building blocks (*pKSB-sgRNA1-3, pJET-5’tan* and *pJET-3’tan*) were cloned by GoldenGate assembly into the destination plasmid *pENTR* together with the building block *pKSB-OpiE2-puro* encoding puromycine resistance, generating the plasmid *pENTR-Tan-OpiE2-puro*. For details on the creation of the *yellow(-)* line see (Klug *et al*., 2022). Primers are given in **Table 1**.

**Table 1:**
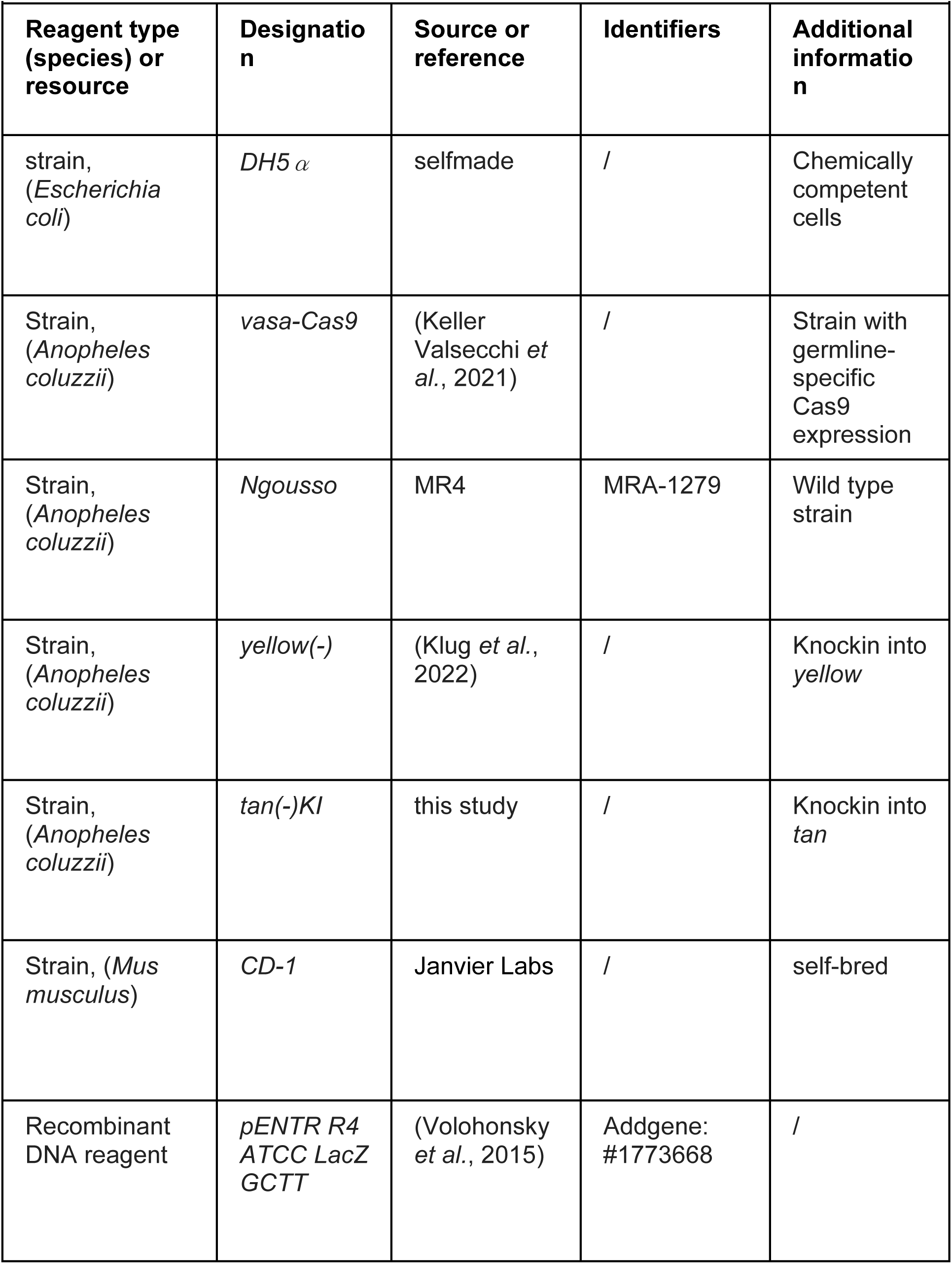
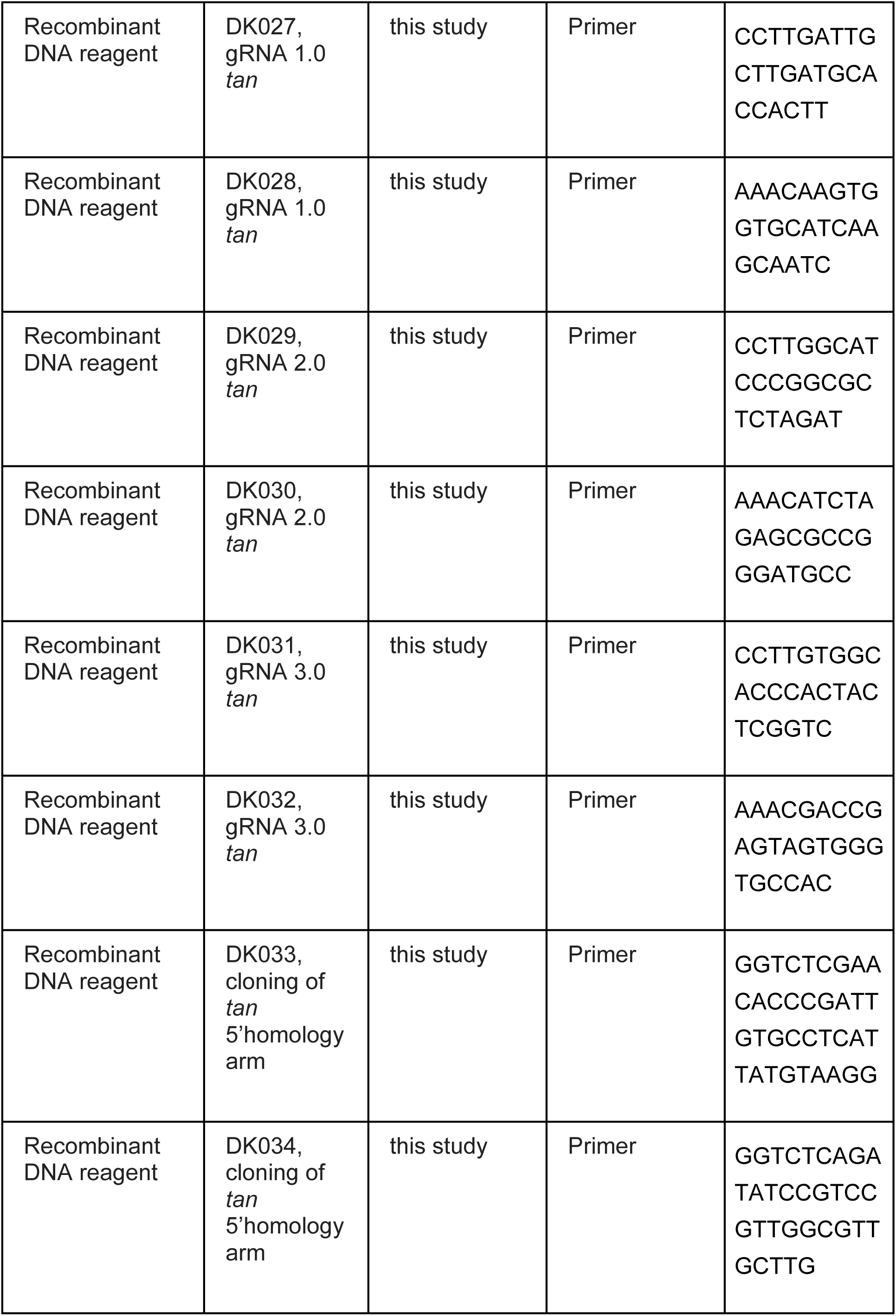
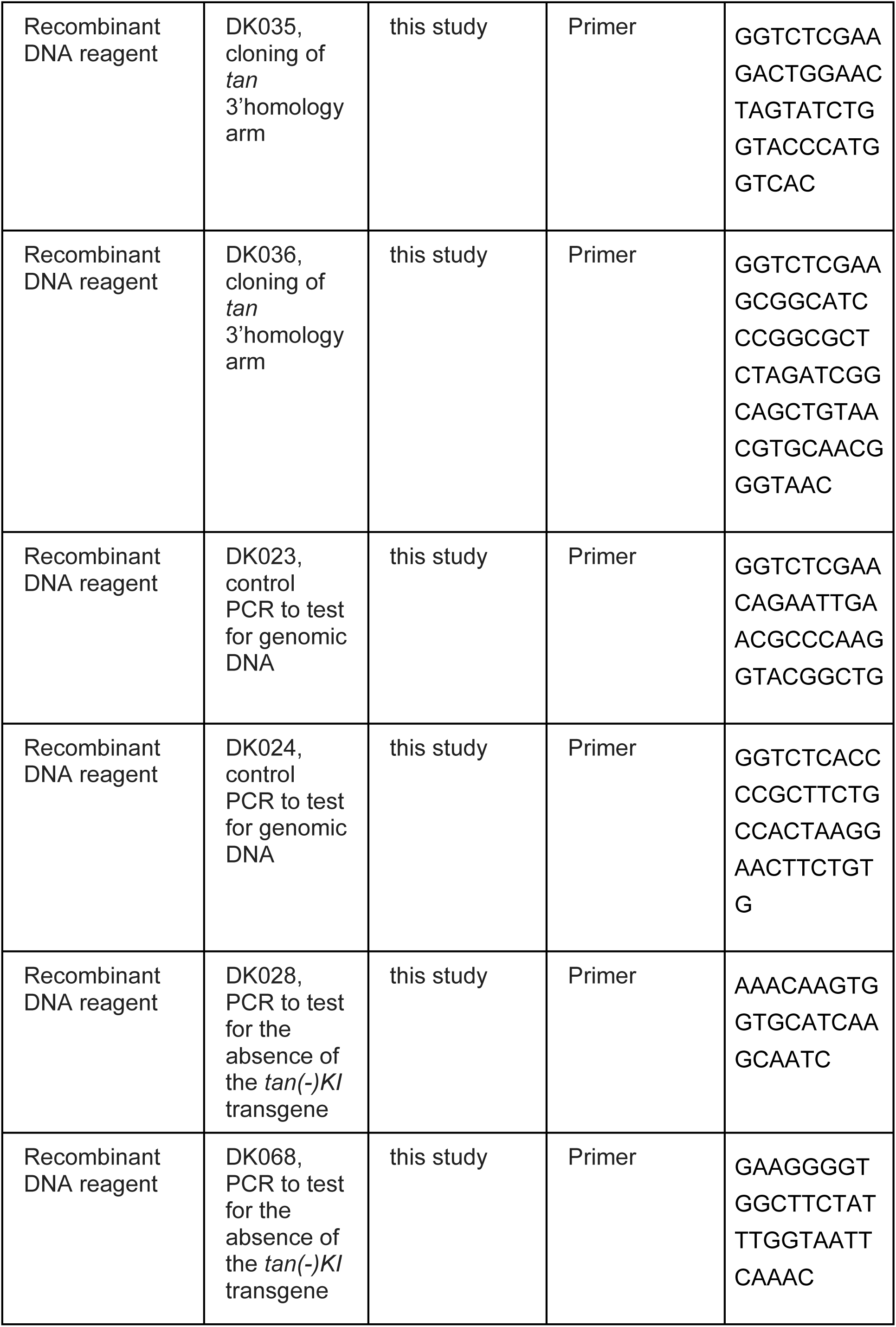
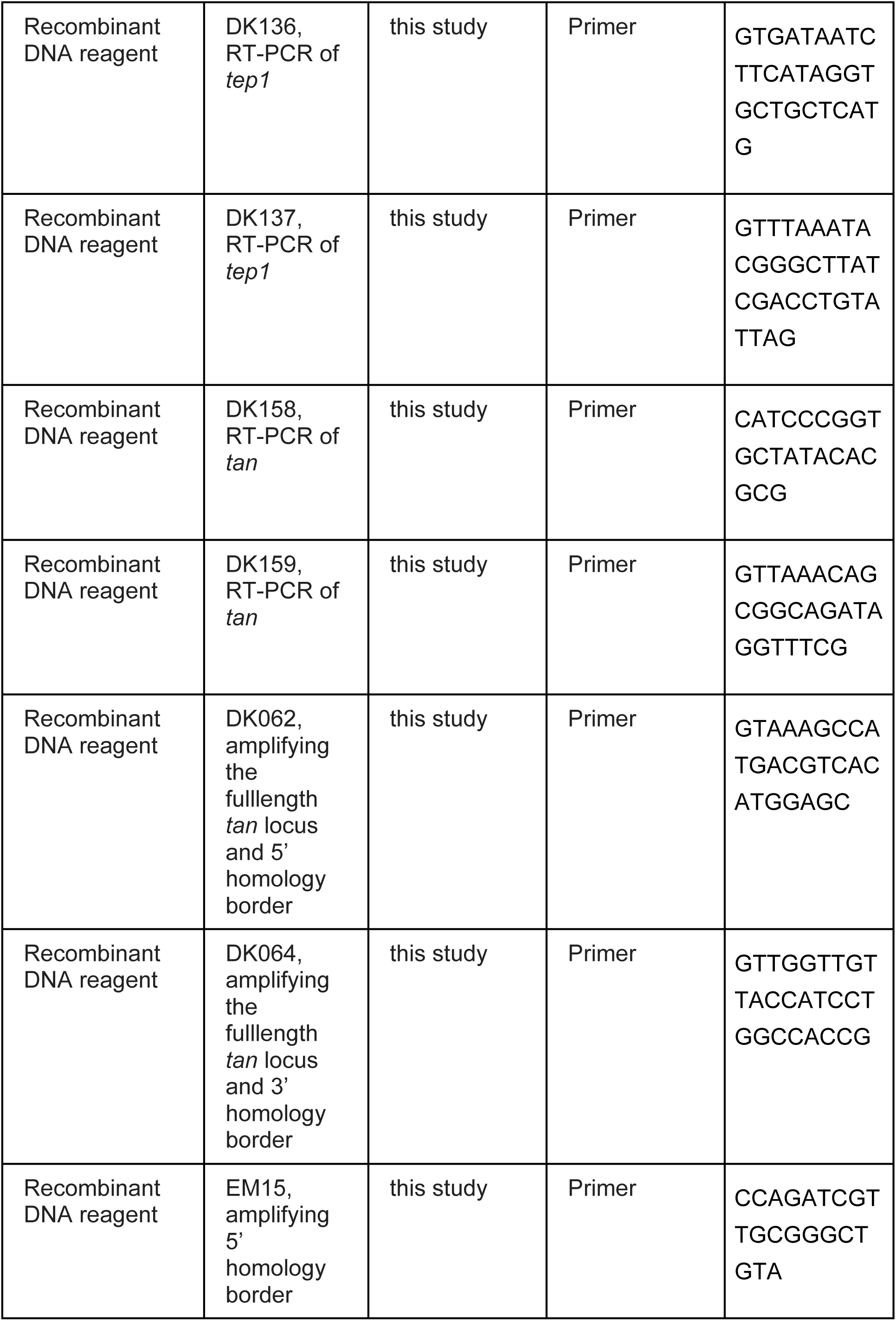
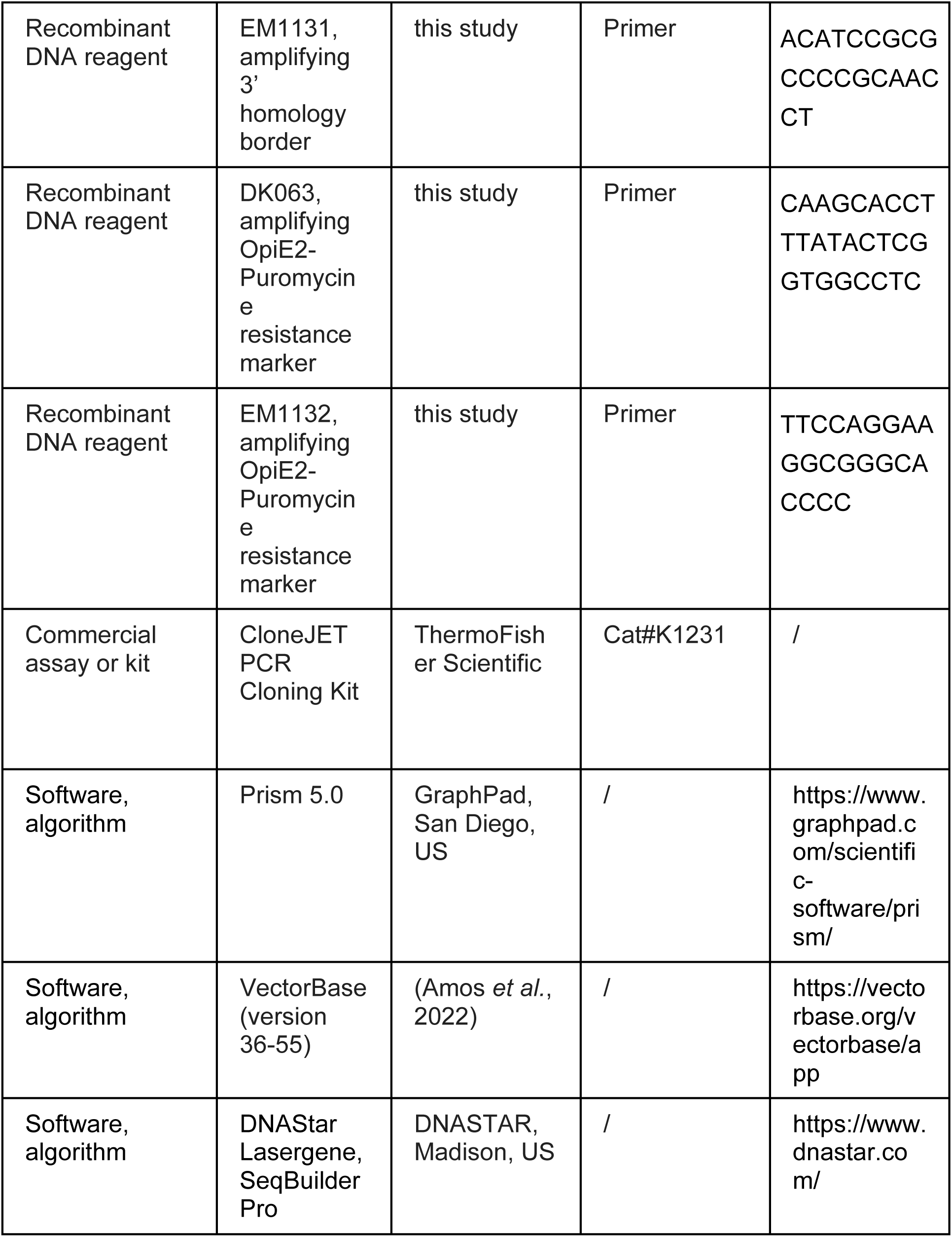
Key Resources.

### Mosquito transgenesis

Freshly laid eggs of the *vasa-Cas9* line were injected as described before (Volohonsky *et al*., 2015). For site-directed knock-in mediated by the Cas9 nuclease, the plasmid *pENTR-Tan-OpiE2-puro* (∼200–300 ng/μL) was diluted in PBS supplemented with 1 μM Scr7 (APExBio, Houston, Texas), an inhibitor of non-homologous end joining. The construct was injected in a batch of approximately 500 eggs obtained from *vasa-Cas9* females. To increase the probablity to generate transgenics and to reduce lethality at the embryo stage, the plasmid was purified using an Endofree kit (Qiagen) before injection. *Yellow(-)* mosquitoes have been generated as described previously (Klug *et al*., 2022)

### Generation of *yellow(-);tan(-)* mosquitoes

Virgin *yellow(-)* females were crossed with *tan(-)* males and the F1 was selfcrossed to allow for recombination events placing both transgenes on the same X chromosome. The F2 offspring was COPAS sorted to isolate EGFP positive individuals which were then treated with puromycin for three days according to standard protocols. Surviving males, which had to be recombinants carrying both transgenes on the same chromosome, were back crossed to *yellow(-)* females. The population was selfcrossed multiple times (≥3 generations) while each generation neonate larvae were treated with puromycine to enrich for the inheritance of the recombinant X chromosome carrying both transgenes. Once lethality of larvae after puromycine treatment was not longer observed, single females were separated in plastic tubes and allowed to lay eggs. This was necessary because one copy of the puromycine resistance marker is sufficient to confer survival after puromycine treatment. After egg laying females were genotyped for the absence of wild type *tan* and the offspring of females homozygous for *yellow(-);tan(-)* was pooled to establish colony.

### Evaluating fitness of *tan(-)* mosquitoes

To estimate the fitness of *tan(-)* mosquitoes I made use of a sexing strain (TS1) inheriting a y-linked transgene expressing GFP (Bernardini *et al*., 2014). By using GFP as marker I was able to separate female and male neonate larvae by COPAS sorting (Marois *et al*., 2012) which made it possible to define the ratio of puromycine resistant females and males in each generation tracing the frequency of the *tan(-)* transgene in each sex over time. In a first step homozygous *tan(-);cas9::RFP+* and *cas9::RFP+* females were crossed separately with TS1 males. Neonate larvae of the F1 of both crosses were COPAS sorted for GFP and RFP expression to isolate males carrying y-linked GFP and one copy of the *cas9::RFP+* transgene. Since the *tan(-)* transgene is X-linked F1 males derived from crossing homozygous *tan(-)* females crossed with TS1 males were carrying the *tan(-)* transgene on their X chromosome making selection with puromycine obsolete. In a second cross [*tan(-);y-GFP;cas::RFP+]^1^*as well as *[y-GFP;cas::RFP+]^1^* males were back crossed either to homozygous *tan(-);cas9::RFP+* or *cas9::RFP+* females. Neonate larvae were COPAS sorted for GFP and homozygous RFP expression. Male larvae of both crosses were raised in synchrony with the *tan(-);cas9::RFP+* and the *cas9::RFP+* colony until adulthood. Pupae of the *tan(-);cas9::RFP+* and *cas9::RFP+* colony were sexed and approximately 75 female pupae of each genotype were hatched in separate cages. To monitor the persistance of the *tan(-)* transgene overtime a cage was set up containing 50 *[y-GFP]^1^;[cas9::RFP+]^2^*males, 50 *[tan(-)]^1^;[y-GFP]^1^;[cas9::RFP+]^2^*males, 50 *[cas9::RFP+]^2^* females and 50 *[tan(-)]^2^;[cas9::RFP+]^2^*females. The mixing of the different genotypes was performed 2-3 days after hatching and all mosquitoes were anaesthetized on ice before being mixed in a cage to prevent a reproductive advantage of any genotype. Females were blood fed 3-4 days after mixing and allowed to lay eggs 48 hours later. Hatched neonate larvae were separated into two batches, either to propagate the colony or to estimate the frequency of the *tan(-)* transgene. To measure transgene frequency 200-250 GFP+ (male) and 200-250 GFP- (female) neonate larvae were sorted in two different 6-cm-diameter petri dishes filled with 8 ml of water. Subsequently dishes containing male or female larvae were treated with a final concentration of 5 ng/µl puromycine (16 µl of a 25 mg/ml stock solution was added) for 3 days (Volohonsky *et al*., 2015). Afterwards survivors were counted and the ratio of puromycine resistant and puromycine susceptible male and female larvae was plotted. The experiment was repeated for further 10 generations to estimate the fitness of *tan(-)* transgene inheriting mosquitoes over time.

### Survival experiments

Four to five day old homozygous *tan(-)* and control females were separated in „survival“ cups and kept at 21°C, 60% humidity and 12 hours light, 12 hours dark cycle. Mosquitoes received 10% sugar solution *ad libitum* provided as soaked cotton pads protected by petri dishes from drying on top of each cup. Cotton pads were moisted every two to three days and completely renewed every seven to ten days. The number of dead mosquitoes was counted every day and dead mosquitoes were immediately removed.

### Evaluating mosquito attraction to UV-B light

To assess the attraction of *tan(-)* mosquitoes to UV-B light behaviour experiments in competition with wild-type mosquitoes were performed. To distinguish between genotypes controls and *tan(-)* mosquitoes were marked with green or orange fluorescent powder (Moon Glow) according to standard protocols (Verhulst, Loonen and Takken, 2013). In brief, 4-10 day old mosquitoes were transferred from breeding cages into paper cups and anaesthetized on ice. In the meantime syringes were filled with either, green or orange fluorescence powder. Once mosquitoes were immobilized cups were transferred into a big plastic bag and mosquitoes were dusted with powder by gently pushing the syringe while holding the needle through the net of the cup. Dusted mosquitoes were shaked a little to distribute the powder equally on all individuals. Subsequently mosquitoes were transferred into fresh cups with sugar pads and allowed to recover overnight under breeding conditions. The color of the powder (orange or green) was alternated between experiments and genotypes to prevent any experimental bias. The next day 20 to 30 mosquitoes of each sex and genotype were transferred into a foldable cage with a volume of 324 l (60 x 60 x 90, Deanyi). Prior to the release of mosquitoes, the cage was prepared with a portable anti-mosquito lamp (Powerology) emitting UV-B light. The lamp attracts mosquitoes by light emitting LEDs that are placed in a circle. Mosquitoes flying through or close to the opening are immediatly sucked into the trap due to the generated underpressure of a fan and subsequently trapped inside. The lamp was placed in the middle of the cage and flanked by two sugar resevoirs which were placed in the shade of the lamp to avoid trapping of feeding mosquitoes. In addition, two normal cotton pads were placed on top of the cage. Mosquitoes which had been released in the cage were allowed to acclimatize for ≥2 hours before the mosquito trap was switched on. Experiments were performed under two different light and collection regims. Initital experiments were performed from 5 pm to 9 am while mosquitoes were kept dark from 5 pm to 2 am and exposed to light from 2 am to 9 am. In this regime mosquito collection was performed the next day at 9 am. In a second setting experiments were performed from 3 pm to 9 am while mosquitoes were kept in the dark from 3 pm to 2 am and exposed to light from 2 am to 9 am. First collection of trapped mosquitos in this setting was performed after two hours (5 pm) and the final collection of trapped mosquitoes was performed at 9 am the next day. Behaviour experiments performed with *yellow(-)* and *yellow(-):tan(-)* mosquitoes were performed in the same way with the exception that dusting with fluorescent powder and recovery overnight was not performed.

### Evaluation of host seeking behaviour and blood feeding success

To assess potential alterations in host-seeking behavior and blood-feeding success, the existing experimental setup previously used to evaluate light attraction was adapted. In the modified configuration, the mosquito trap was removed, while sugar feeders remained in place. In each experiment, 20 control (wild-type or *yellow(-)*) and 20 *yellow(-);tan(-)* female mosquitoes (4–10 days post-eclosion) were released into a cage and allowed to acclimate for at least 2 hours. Following acclimation, a human arm was introduced as a host stimulus through a small opening at the bottom of the cage. Mosquitoes were allowed to bite for 15 minutes. After the exposure period, the cage was carefully folded and transferred to a −20 °C freezer for 10 minutes to immobilize the mosquitoes. Subsequently, all individuals were collected, and the number of blood-fed females was recorded.

### Software

Graphs and statistical analyses were generated using GraphPad Prism 5.0 (GraphPad, San Diego, CA, USA). Figures and illustrations were created with Adobe Illustrator CS5.1 (Adobe, San Jose, CA, USA). ChatGPT (GPT-4o, OpenAI, San Francisco, CA, USA) was used to assist in refining the manuscript’s phrasing, spelling, and grammar.

## Declarations

### Ethics approval for animal experiments

All experiments were conducted in accordance with Directive 2010/63/EU of the European Parliament, which governs the protection of animals used for scientific purposes. Our animal facility is accredited under license number I-67-482-2, issued by the veterinary authorities of the Bas-Rhin département (Direction Départementale de la Protection des Populations). The use of animals in this study, as well as the generation and utilization of transgenic organisms (bacteria and mosquitoes), was authorized by the French Ministry of Higher Education, Research and Innovation under authorization numbers APAFIS#20562–2019050313288887 v3 and 3243, respectively.

### Consent to participate

Not applicable.

### Consent for publication

Not applicable.

### Availability of data and materials

All data have been made available within this manuscript. Materials are available from the corresponding author on reasonable request.

### Authors’ contributions

DK designed the study, performed experiments, curated and analyzed the data, made the figures and wrote the manuscript.

## Acknowledgements

We thank Nathalie Schallon and Amandine Gautier for their support in rearing *Anopheles coluzzii* mosquitoes, and Pia Thiele for technical assistance with experiments. I am also grateful to the Mosquito Immune Responses (MIR) team for helpful discussions and weekend support with mosquito maintenance. Special thanks go to Eric Marois for his contribution to mosquito transgenesis, and to Stéphanie Blandin for her continuous support and guidance in experimental planning. I am also grateful to Dr. Joerg Klug for his careful review and insightful comments on the manuscript.

---

**Figure S1.**
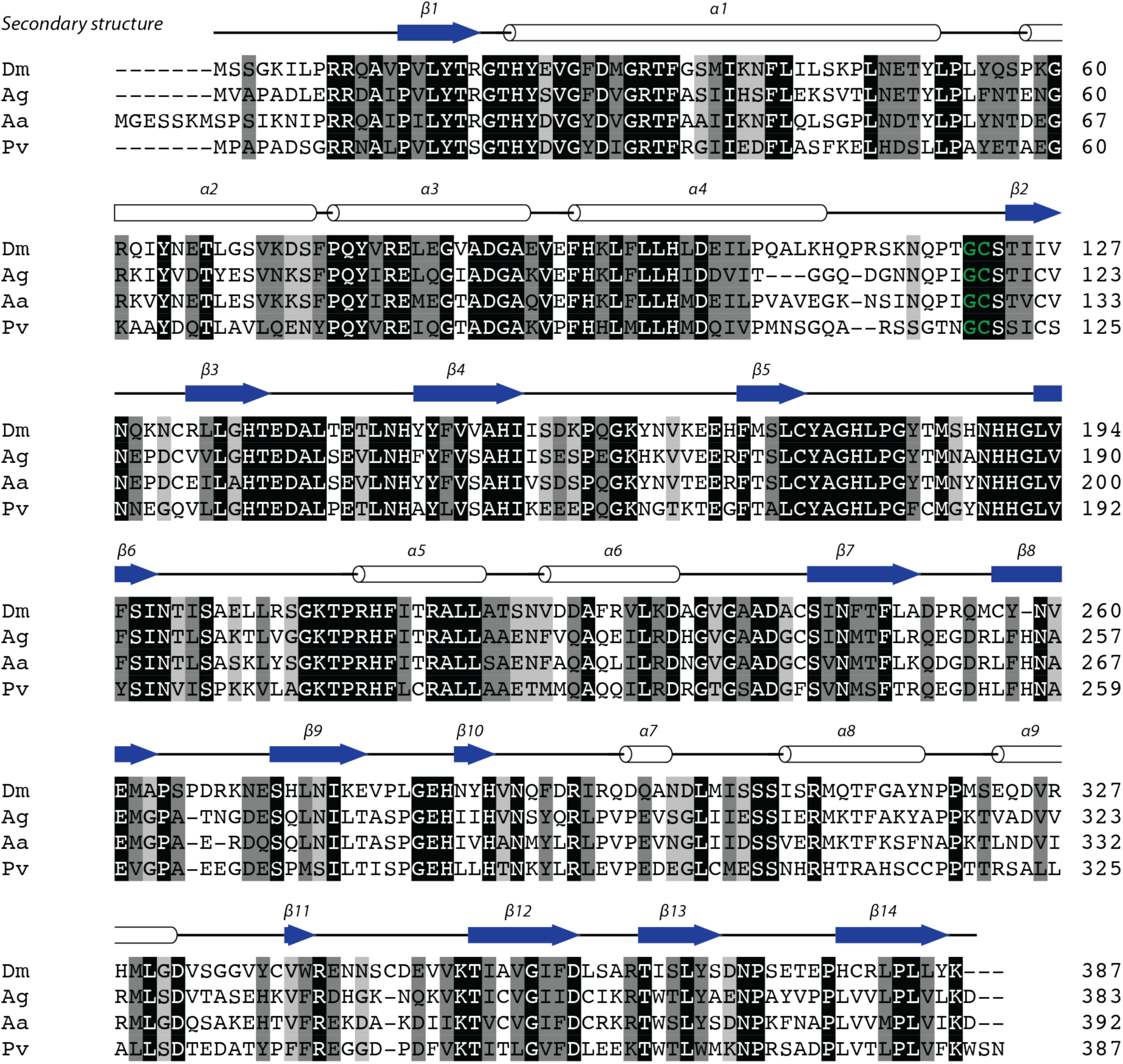
Tan is highly conserved in insects and crustaceans. Multiple sequence alignment of Tan homologues from the fruity fly *Drosophila melanogaster* (Fbgn0086367), the malaria mosquito *Anopheles gambiae* (AGAP000023), the yellow fever mosquito *Aedes aegypti* (AAEL023223) and the whiteleged shrimp *Penaeus vannamei* (LOC113823002). The secondary structure predicted with i-TASSER based on the sequence of AgTAN is indicated above the alignment (arrows indicate beta-strands, whereas barrels indicate alpha-helices). Residues that are conserved are written in white and highlighted in black, mostly conserved residues are highlighted in grey. The cleavage site (Gly-Cys motif) between the small and the big subunit is highlighted in green.

**Figure S2.**
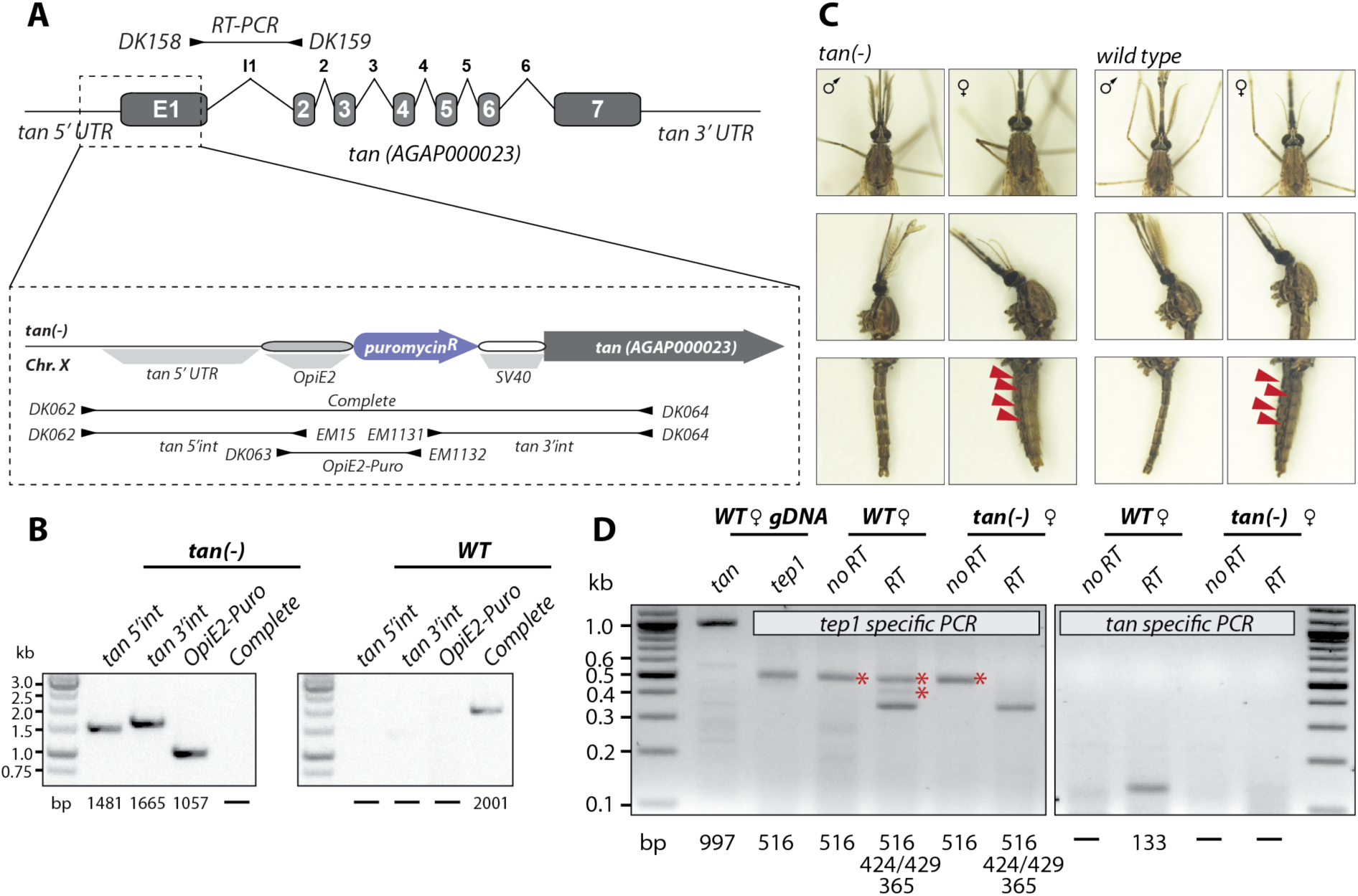
Generation and characterization of the *tan(-)* mosquito line. **A)** Scheme illustrating the generation of *tan(-)* mosquitoes. Transgenic mosquitoes were generated by Cas9-mediated DNA cleavage and repair using a provided template carrying a puromycin resistance cassette to replace the first half of exon 1 of the *tan* gene (AGAP000023), including the start codon. The transgene consists of the puromycin N-acetyl transferase resistance gene flanked by the viral *Op*IE2 promoter and an SV40 terminator (Volohonsky *et al*., 2015). Primers used for genotyping and amplified PCR products are indicated with arrowheads and black lines below the scheme, respectively. Note that the illustration is not drawn to scale. **B)** Genotyping of the *tan(-)* colony. Expected fragment lengths for the 4 PCRs are indicated below the gel, and marker sizes on the left of the gel. Note that the complete PCR fragment (DK062/DK064) could not be amplified from *tan(-)* mosquitoes because of the large fragment length after integration of the resistance cassette. **C)** Adult female and male *tan(-)* mosquitoes in comparison to wild-type (*Ngousso*). Shown specimens have an age of 4-7 days post emergence. Colonies of both lines were bred in synchrony to avoid pigment alterations due to age-dependent changes in cuticle pigmentation. Red arrowheads point toward less pigmentated areas in the abdomen of *tan(-)* females. **D)** The absence of *tan* transcription in the *tan(-)* line was verified by RT-PCR on cDNA samples generated from wild type (wt) and *tan(-)* females. Primers amplifying a fragment of *tep1* (exon #1 to exon #3) were used as positive control. Red asterisks highlight PCR products matching with the expected length of unspliced pre-mRNA or genomic DNA as well as partially spliced mRNA.

**Figure S3.**
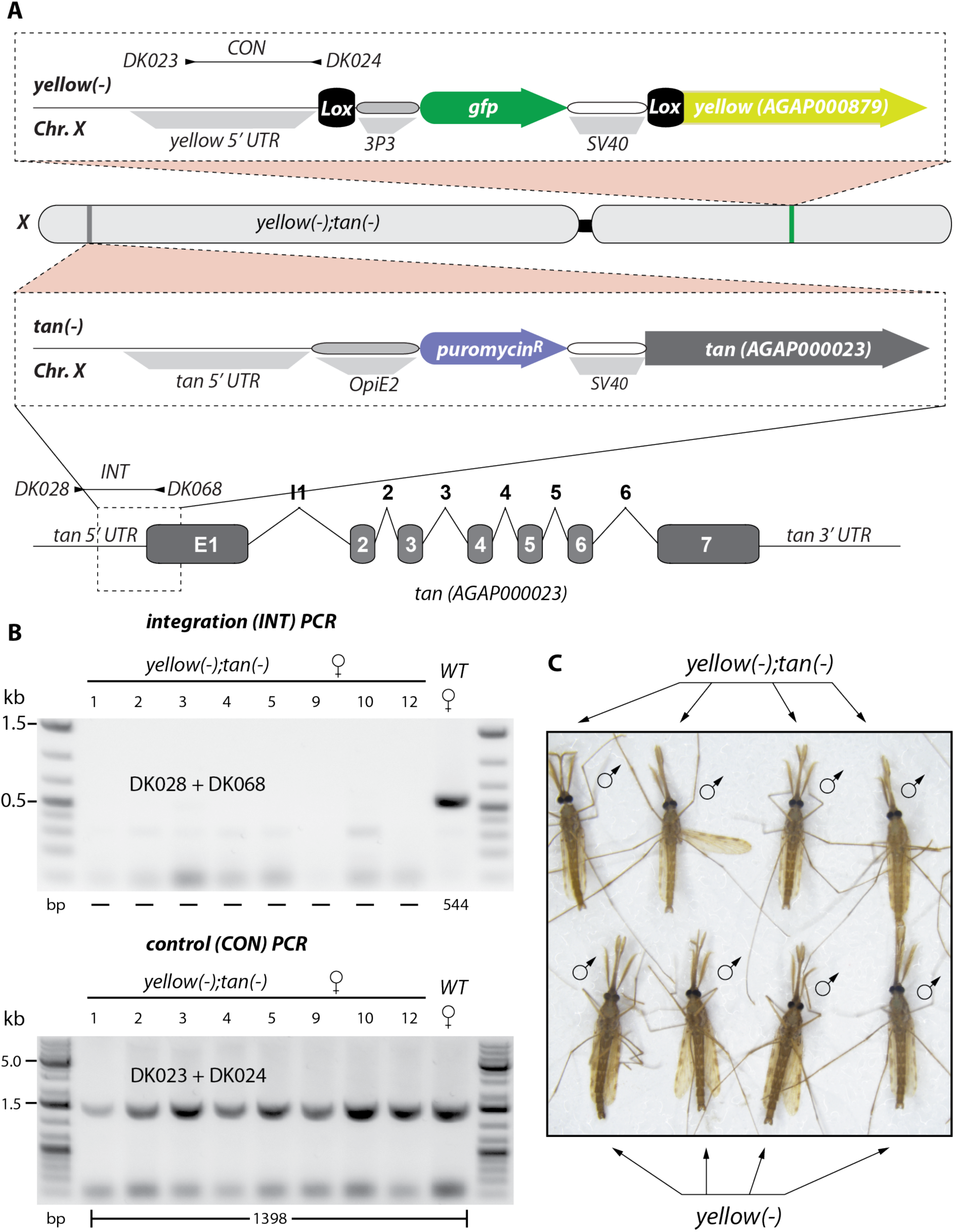
Validation and characterisation of *yellow(-);tan(-)* mosquitoes. **A)** Scheme illustrating the expected genotype of *yellow(-);tan(-)* mosquitoes. Note that both transgenes reside on chromosome X. Genotyping primers and amplified PCR products are indicated by black arrows and black lines, respectively. **B)** Genotyping PCRs of eight females homozygous for *yellow(-);tan(-)*. The top INT PCR amplifies a fragment between the 5’UTR and the first exon of *tan* which is only present in wild type but absent in all eight females confirming the homozygosity for *yellow(-);tan(-)*. The bottom CON PCR amplifies the 5’UTR of *yellow* present in wild type but also in *yellow(-)* mosquitoes. It serves as control for the presence of genomic DNA in all samples. Homozygous *yellow(-)* neonate mosquito larvae had been sorted in advance based on their EGFP fluorescence. **C)** Image of four *yellow(-);tan(-)* males (top row) and four *yellow(-)* males (bottom row). Note the brightening of the cuticle in males carrying both transgenes in comparison to males lacking expression of *yellow* only

**Figure S4.**
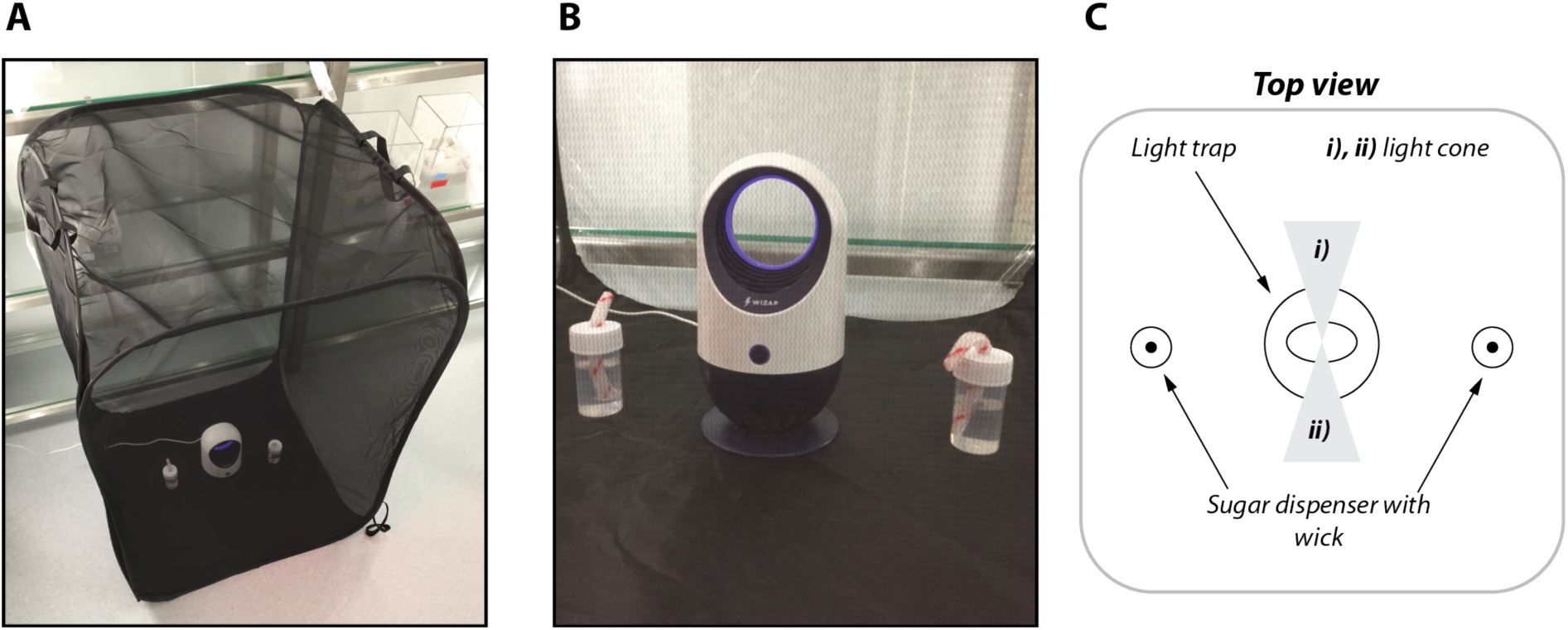
Experimental setup to evaluate attraction of malaria mosquitoes to UV-B light. **A)** Cage setup under laboratory conditions. **B)** Placement of light trap and sugar dispensers within the cage. **C)** Illustration of the experimental setup in top view. Note that sugar dispensers were placed out of light cones emitted by the trap to avoid the catching of sugar feeding mosquitoes.

**Figure S5.**
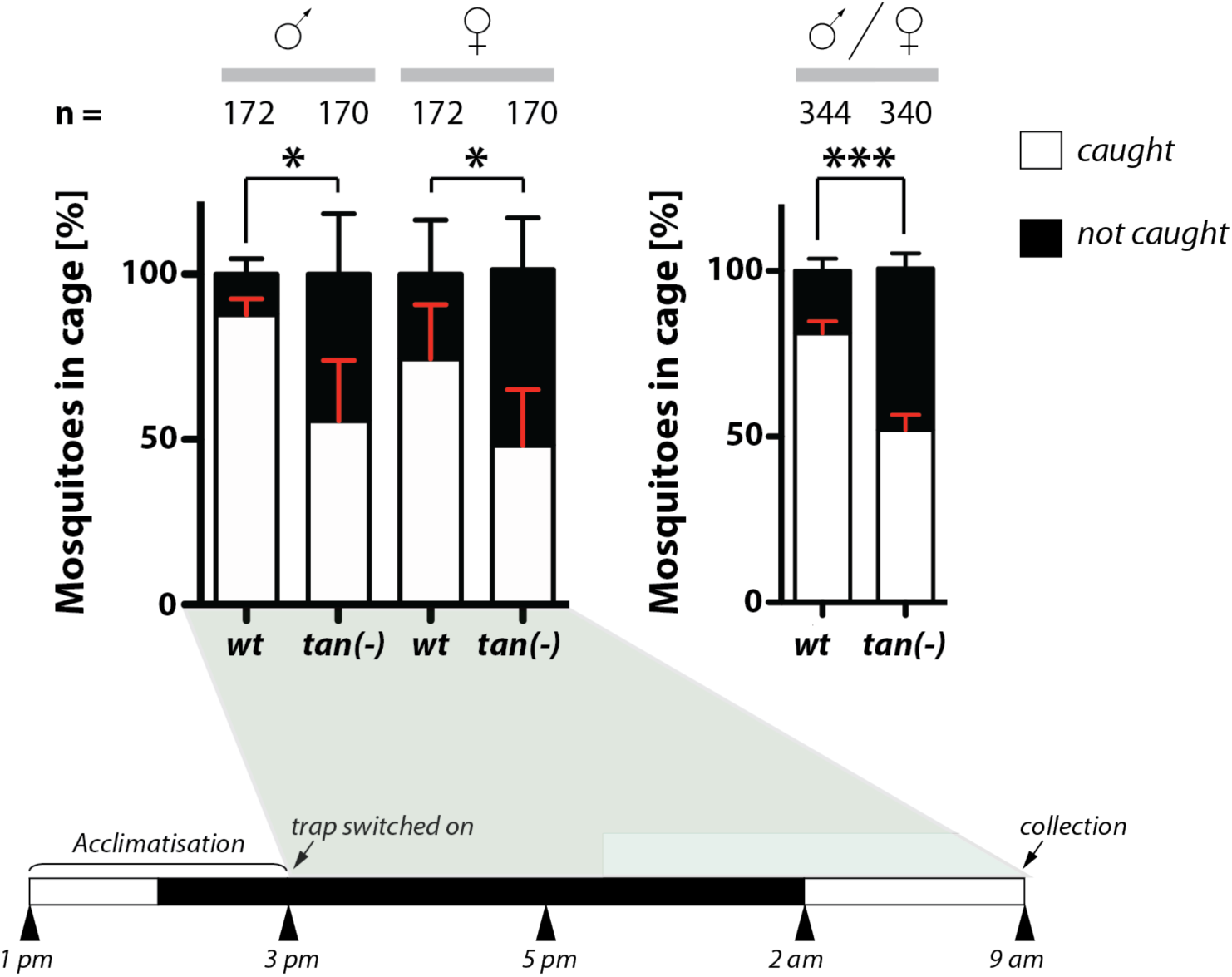
Combined data of UV-B catching experiments performed with *tan(-)* and *yellow(-);tan(-)* mosquitoes in comparison to wild-type. Graphs display the dombined data of Figure 3D and and Figure 4C of the *tan(-)* and the *yellow(-);tan(-)* line in comparison to wild-type. The left graph displays the sex-specific catching rates while the graph on the right combines catching rates of males and females for each genotype. Numbers above each column indicate the number of tested mosquitoes. Datasets were tested for significance with Mann Whitney test. p value = 0.0101* (males), p value = 0.0150* (females), p value = 0.0004*** (both sexes combined).

